## Supplemental text S1 for "New alignment-based sequence extraction software (ALiBaSeq) and its utility for deep level phylogenetics"

Supplemental text S1. Details of the plant ODB SCO dataset preparation, assessment methods, and results.

**Methods**

**Samples**

To verify the performance of ALiBaSeq on a pair of divergent plant species, we used *Arabidopsis thaliana* (L.) Heynh. and *Amborella trichopoda* Baill., both of which have annotated genomes available. TAIR10 genome assembly, coding sequences, and proteins for *A. thaliana* and AMTR1.0 genome assembly, coding sequences, and proteins for *A. trichopoda* were obtained through the Ensembl project. We chose *Arabidopsis* to represent a bait species and *Amborella* to be a sample which sequences were to be retrieved. Thus, in addition to the AMTR1.0 scaffolds, based on this assembly we simulated reads with 40x depth using Art as described in the material and methods.

**Baits**

Bait sequences were prepared from *Arabidopsis* generally following the ODB SCO section of the bait methods in the main text. We used OrthoDB v9 to retrieve the single copy orthologs shared between *Arabidopsis* and *Amborella*, filtered the sequences to have at least 100AA and no more 10% gaps, and compiled both CDS and protein bait sequences, resulting in 3655 loci. We then selected first 1000 loci for the evaluation purposes. The resulting dataset had locus lengths ranging from 300 to 14,085 bp, and nucleotide pairwise sequence distances ranging from 7.31% to 53.03% (average 34.01%) and 2.12% to 67.61% (average 35.59%) on the protein level.

**Software and settings**

We ran the best performing ALiBaSeq configurations (dc-megablast and tblastx) with both strict and relaxed RBH options, phyluce, and HybPiper. HybPiper’s performance with relaxed search settings was on par with the default settings (as opposed to the insect ODB SCO dataset), and relaxed search results were thus used. Assessment methods follow the methods used for the insect ODB SCO dataset.

**Results (Fig. S1)**

The plant ODB SCO dataset proved to be more difficult, in part due to the larger average pairwise sequence distance between the bait and the sample. On the original assembly ALiBaSeq found 750-870 loci depending on the settings. On the 40x data, ALiBaSeq with the relaxed RBH check outperformed HybPiper by finding ~900 loci out of 1000 and having twice less false positives.
