## Supplementary figures and images for "New alignment-based sequence extraction software (ALiBaSeq) and its utility for deep level phylogenetics"

### Figure S1

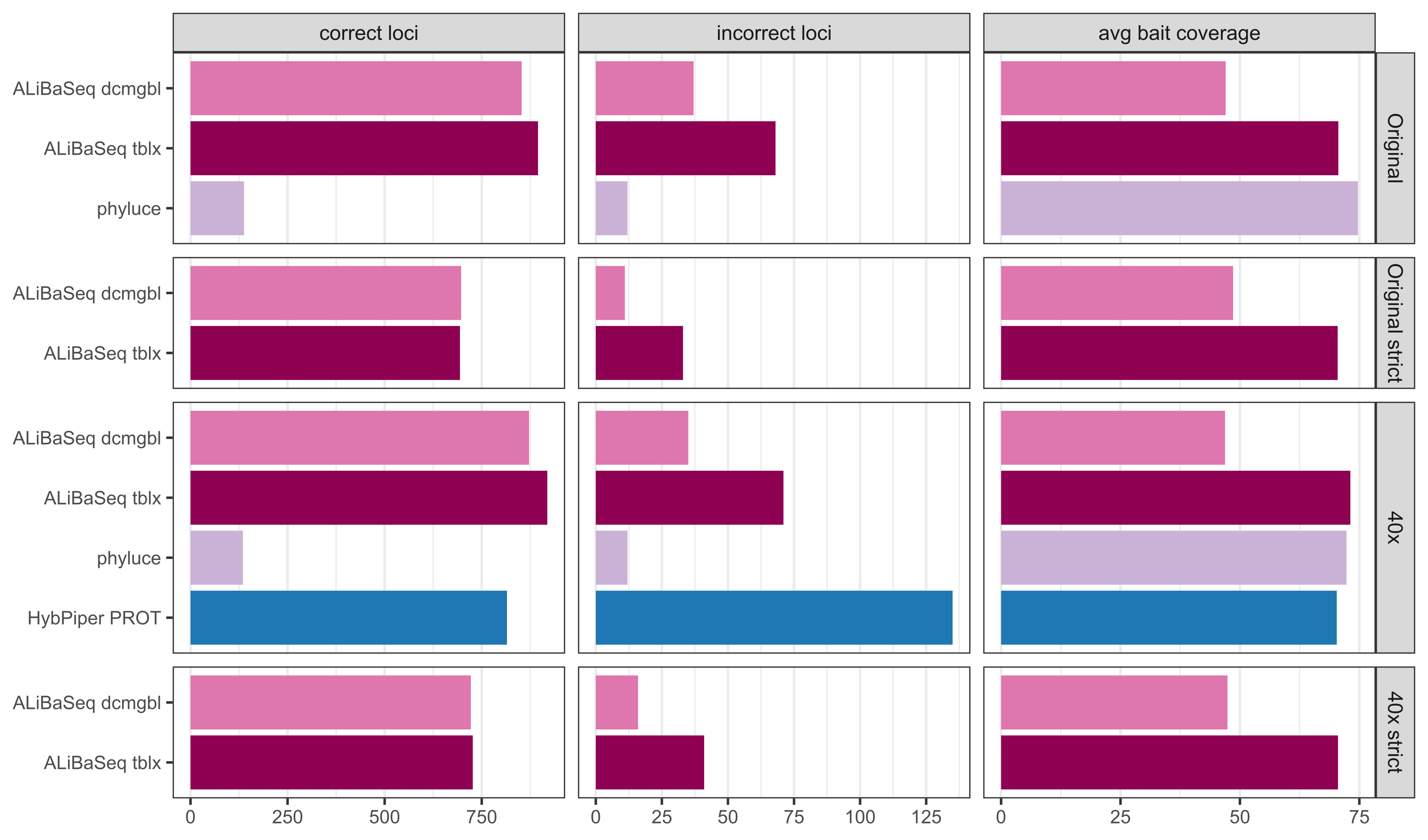
