## Supplementary material for "New alignment-based sequence extraction software (ALiBaSeq) and its utility for deep level phylogenetics": Table S2

Table S1. Program versions and settings used for evaluations.

| Software | Version / Git repo date | Options |
| --- | --- | --- |
| ALiBaSeq | V1.1 | Similarity searches: blastn, blastn with word size set to 9, discontinuous megablast, tblastn, tblastx, nhmmer, and phmmer   - BLAST: -evalue 1e-03 -num_threads 32 - HMMER: --cpu 32 -E 1e-03   RBH similarity searches: discontinuous megablast with settings as above  General UCE analysis: -f S -x s -e 1e-03 -c 1 -i 50 -m b-i-e  General ODB analysis: -f S -x a -e 1e-03 -c 1 --is -i 50 --ctg-ovlp 30 --hit-ovlp 50 --amalgamate-hits -m b-i-e  Original assembly ODB analysis: -m b/e-I instead of -m b-i-e  blastn- and megablast-based analysis: --ac dna-dna  tblastx-based analysis: --ac tdna-tdna  tblastn-based analysis: --ac aa-tdna  nhmmer-based analysis: --ac dna-dna --bt hmmer15  phmmer-based analysis: --ac aa-tdna --bt hmmer22  Additional parameter used to turn on the strict RBH check: --rm-rec-not-found |
| Assexon | Thu Aug 15 07:37:41 2019 | the assembly-based version: --cpu 32 --E 1e-03 --min_len 50  the read-based version: --cpu 32 --E_value_ubandpx 1e-03 --E_value_reblast 1e-03 --similarity 50 |
| aTRAM | Fri Sep 13 17:01:05 2019 | atram_preprocessor.py was run with default parameters  DNA-based aTRAM parameters: --cpus 32 -a spades --max-memory 120 --evalue 1e-03 --bit-score 0 --contig-length 30  Protein-based aTRAM parameters: --cpus 32 -p -a velvet --max-memory 120 --evalue 1e-03 --bit-score 0 --contig-length 30  Exon-stitching was run with default parameters |
| FortyTwo | v 0.190820 | yaml-generator-42.pl parameters: --run_mode=phylogenomic --evalue=1e-03 --homologues_seg=yes --max_target_seqs=10000 --templates_seg=no --ref_brh_mode=on --ref_org_mul=1.0 --ref_score_mul=0.99 --trimming_mode=off --ls_action=keep --aligner_mode=blast --ali_patch_mode=off --ali_cover_mul=1.1 --tax_reports=on |
| HybPiper | v 1.2 Wed Aug 8 09:43:03 2018 | DNA-based reads_first.py parameters: --cpu 32 --bwa --evalue 1e-03 --thresh 50 --length_pct 50 --depth_multiplier 1  General protein-based reads_first.py parameters: --cpu 32 --evalue 1e-03 --thresh 50 --length_pct 50 --depth_multiplier 1  Protein-based reads_first.py parameters for ODB search on 20x and 40x reads: --cpu 32 --thresh 65 --length_pct 90 --depth_multiplier 10 |
| Kollector | Wed Mar 21 19:16:14 2018 | -j 32 -r 0.5 |
| Phyluce | v 1.6.7 | Generally, samples were processed according to phyluce tutorial 3 (but see main text) with the following parameters:   - phyluce_probe_run_multiple_lastzs_sqlite: --identity 50 --coverage 50 - phyluce_probe_slice_sequence_from_genomes: --flank 0 - phyluce_assembly_match_contigs_to_probes --min-identity 50 --min-coverage 50 |
| Trimmomatic | 0.36 | PE LEADING:25 TRAILING:25 SLIDINGWINDOW:4:25 MINLEN:36 |
| SPAdes | 3.12.0 | -t 32 -m 380 |
| Art | MountRainier | General parameters: -ss HSXt --len 150 --mflen 225 --sdev 75  For different depths the parameter -f was adjusted to 1, 3, 5, 10, 20, and 40 |
| Mafft | v7.427 | UCE dataset parameters: --thread 32 --inputorder --globalpair --maxiterate 1000  ODB dataset parameters: --thread 32 --inputorder --genafpair --maxiterate 1000 |
